# MMMVI: Detecting SARS-CoV-2 Variants of Concern in Metagenomic Wastewater Samples

**DOI:** 10.1101/2021.06.14.448421

**Authors:** Dillon O.R. Barker, Cody J. Buchanan, Chrystal Landgraff, Eduardo N. Taboada

## Abstract

**Motivation:** SARS-CoV-2 is the causative agent of the COVID-19 pandemic. Variants of Concern (VOCs) and Variants of Interest (VOIs) are lineages that represent a greater risk to public health, and can be differentiated from the wildtype virus based on unique profiles of signature mutations. Metagenomic sequence analysis of wastewater represents an emerging form of surveillance that can capture early signal for these variants in a community prior to detection through public health testing or genomic surveillance activities. However, because multiple viral genomes are likely to be present in a metagenomic sample, additional analytical scrutiny of the sequencing reads beyond variant calling is needed to increase confidence in diagnostic determinations.

**Results:** Where multiple signature mutations are present on a given read, they can be used as enhanced biomarkers to confirm the presence of a VOC/VOI in the sample. We have developed MMMVI, a tool to aggregate and report on the likely presence of a VOC/VOI in a set of metagenomic reads based on the detection of reads bearing multiple signature mutations.

**Availability:** MMMVI is implemented in Python, and is available under the MIT licence from https://github.com/dorbarker/voc-identify/

## 1 Introduction

Wastewater testing for SARS-CoV-2 can offer several advantages that can help address testing gaps and complement more traditional forms of laboratory-based surveillance. Wastewater surveillance is non-invasive, can capture signal from asymptomatic and positive cases not detected via the public health system, and can detect early-warning data predictive of future clinical trends (Medema *et al*., 2020; D’Aoust *et al*., 2021). Metagenomic sequencing provides a mechanism to characterize the genetic diversity of SARS-CoV-2 present in wastewater, and presents a snapshot of the lineages circulating within the community. Importantly, these data can be screened for the presence of SARS-CoV-2 Variants of Concern (VOCs) and Variants of Interest (VOIs) based on the profile of signature mutations observed in the sample (Hadfield *et al*., 2018; Rambaut *et al*., 2020; Public Health England, 2021). However, a sample may not contain the complete set of mutations corresponding to any single VOC/VOI. This could be due to incomplete sequencing of the genome, or indicative of the early introduction, or localized outbreak of one or more VOCs/VOIs in the community. It is also possible to observe signature mutations from multiple VOCs/VOIs in a single sample, and some signature mutations (*e*.*g*., N501Y) have emerged independently in multiple VOCs/VOIs. Thus, the criteria (*i*.*e*., the number and/or set of specific signature mutations required) to stipulate the presence of a VOC/VOI in a metagenomics context are not well defined.

We propose an approach that takes advantage of signature mutations that occur in close proximity to one another in the SARS-CoV-2 genome. If we assume that an individual sequencing read is ultimately derived from a distinct viral genome, then any mutations identified on that read should also be derived from that viral genome. Therefore, the combination of signature mutations found on a single read can be used as an enhanced biomarker that is more sensitive than variant calling for the detection of a VOC/VOI, but also suitable as a surrogate marker in samples where the genome is incompletely sequenced.

Here, we present MMMVI, a command line tool that automates the categorisation and quantification of reads based on all combinations of signature mutations they encompass. It generates a number of detailed reports to more confidently inform the presence of, or differentiate between, one or more VOCs/VOIs in a sample. MMMVI is compatible with data generated from both Illumina and Oxford Nanopore Technologies platforms, and can be readily run both on consumer-grade hardware and in parallel on a computer cluster.

## 2 Description

### 2.1 Data

A wastewater sample positive by RT-qPCR detection for the B.1.1.7/N501Y.V1/Alpha SARS-CoV-2 VOC was amplified using a tiled-PCR amplification method and sequenced on both the Illumina MiSeq and Oxford Nanopore Technologies MinION platforms (BioProject: PRJNA708265). Reads from each sequencing run were independently mapped to the reference genome, SARS-CoV-2 isolate Wuhan-Hu-1 (MN908947.3) (Wu *et al*., 2020).

### 2.2 Input

MMMVI takes three files as input: the sample in BAM format aligned to the reference genome (Li *et al*., 2009), the complete reference genome in FASTA format, and variant definitions. These definitions can be provided either as a tabular text file matching the format described in Table 1, or as YAML files matching the Public Health England Standardised Variant Definitions specification (Public Health England, 2021).

**Table 1.** Example of the tabular mutation file format required by MMMVI.

| VOC/VOI | Mutation |
| --- | --- |
| B.1.1.7 | C23604A |
| B.1.1.7 | GAT28280CTA |
| B.1.1.7 | [21765-21770]del |
| P.1 | 28262AACA |

**Table 2.**
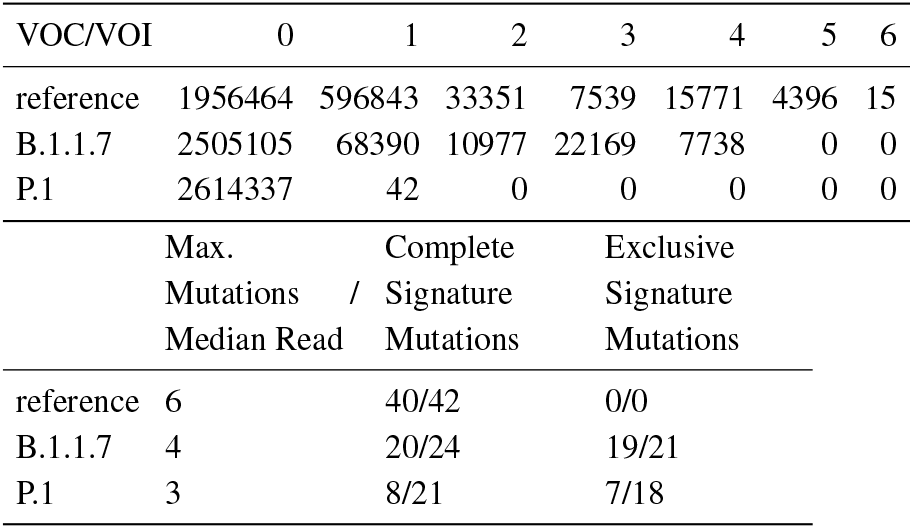
An example of the summary table produced by MMMVI from Illumina sequencing of a metagenomic wastewater sample.

The first line in Table 1 describes substitution of C in the wildtype by A in B.1.1.7 at reference genome position 23604 in the sense strand. Similarly, the second line shows a three base substitution of GAT by CTA, beginning at position 28280. The third line is a 6 base deletion relative to the reference beginning at position 21765. On the fourth, a four base insertion between reference positions 28262 and 28263 is indicated. All positions are 1-based, *i*.*e*. position 1 is the first base in the reference genome. All intervals are closed.

### 2.3 Mutation Identification

MMMVI makes use of pandas and pysam (https://github.com/pysam-developers/pysam) (McKinney *et al*., 2011; PySAM Developers, 2019) for efficient processing.

Each read is compared to the set of mutations provided by the mutations table. If a read spans all the reference positions for a given mutation, its sequence at those positions is compared with the mutation. If the read contains all of the substitutions or deletions described by the mutation, the read is considered to have that mutation. Gaps in the read alignment are considered deletions. A read may contain multiple mutations, and each unique combination of mutations is considered a read species.

### 2.4 Output

All outputs are returned to the user as delimited text files, with the delimiter optionally selectable by the user.

#### 2.4.1 Summary Table

The summary file contains a table indicating the count of reads possessing *N* mutations for each VOC/VOI, as well as the wildtype. The numerical column headers indicate *N* mutations found on the read, and the left column shows the variant names. The table values for these show the count of reads with *N* mutations for each variant. The maximum mutations per median read indicates the greatest theoretical number of mutations for a given VOC/VOI that can fit on a single read based on the median read length calculated for the sample. Also shown are the fraction of signature mutations that are represented in the sample, as well the fraction of signature mutations which are exclusive to that VOC/VOI.

#### 2.4.2 Co-occurrence Matrices

The co-occurrence matrices show the frequencies of cases where multiple mutations are present on a given read.

These matrices are given both as absolute counts and as relative counts. The relative count expresses, for each pair of mutations *X* and *Y*, the count of reads exhibiting both *X* and *Y* as a proportion of the count of all reads possessing *X*.

#### 2.4.3 Read Report

This report shows every sequencing read given as input and alleles present for each VOC/VOI and the wildtype. The alleles are expressed as genomic positions, with coordinates placed in the column or columns which correspond to a VOC or the wildtype.

#### 2.4.4 Read Species Report

The read species report shows every unique combination of mutations found on a single read, as well as the positions that are mutated, the nucleotide or INDEL present at those positions, and the number of times a given combination was observed in the dataset.

Further, it provides a bit array for each VOC/VOI and the wildtype, indicating at each position which variant shares the observed nucleotide or INDEL. Finally, each combination is expressed as a proportion relative to the number of reads spanning the relevant positions, and as a proportion relative to the total number of reads in the dataset.

## 3 Conclusion

MMMVI uses a command line interface for user interaction. For ease of automation, we also provide a Snakemake workflow for executing MMMVI jobs (Mölder *et al*., 2021).

Table 3 shows the performance of MMMVI on inputs generated using the Illumina MiSeq and Oxford Nanopore Technologies MinION. We searched for eleven VOCs or VOIs plus wildtype, comprising 153 variant positions.

**Table 3.** Computing resource consumption on example inputs when searching for eleven variants and the wildtype.

| Instrument | Read Count | Elapsed (min) | Memory Usage (GB) |
| --- | --- | --- | --- |
| MiSeq | 2 614 393 | 44 | 9.32 |
| MinION | 84 078 | 8 | 0.48 |

MMMVI facilitates detection of SARS-CoV-2 VOCs in metagenomic sequence data sourced from wastewater. Although it was designed for use with wastewater samples, we believe this approach is applicable to other sample types.

An example of results generated by MMMVI using Illumina MiSeq and Oxford Nanopore Technologies MinION data from BioProject PRJNA708265 are provided as Supplemental Data.

## Supporting information

Example Output

## Acknowledgements

We would like to thank Robert Delatolla, Tyson Graber and Lawrence Goodridge for organising the collection of and providing access to the wastewater sample used for this analysis.

